# Interplay of mesoscale physics and agent-like behaviors in the parallel evolution of aggregative multicellularity

**DOI:** 10.1101/2020.06.03.133025

**Authors:** Juan A. Arias Del Angel, Vidyanand Nanjundiah, Mariana Benítez, Stuart A. Newman

## Abstract

Myxobacteria and dictyostelids are prokaryotic and eukaryotic multicellular lineages, respectively, that after nutrient depletion aggregate and develop into structures called fruiting bodies. The developmental processes and the resulting morphological outcomes resemble one another to a remarkable extent despite their independent origins, the evolutionary distance between them and the lack of traceable levels of homology in the molecular mechanisms of the groups. We hypothesize that the morphological parallelism between the two lineages arises as the consequence of the interplay, within multicellular aggregates, between *generic processes*, physical and physicochemical processes operating similarly in living and non-living matter at the mesoscale (~10^-3^-10^-1^ m) and *agent-like behaviors*, unique to living systems, characteristic of the constituent cells. To this effect, we analyze the relative contribution of the generic and agent-like determinants in the main phenomena of myxobacteria and dictyostelid development, and their roles in the emergence of their shared traits. We show that as a consequence of aggregation collective cell-cell contacts mediate the emergence of liquid-like properties, making nascent multicellular masses subject to new sets of patterning and morphogenetic processes. In both lineages, this leads to behaviors such as streaming, rippling, and rounding up, similar to effects observed in non-living fluids. Later the aggregates solidify, leading them to exhibit additional generic properties and motifs. We consider evidence that the morphological phenotypes of the multicellular masses deviate from the predictions of generic physics due to the contribution of agent-like behaviors. These include directed migration, quiescence, and oscillatory signal transduction of the cells mediated by responses to external cues acting through species-specific regulatory and signaling mechanisms reflecting the evolutionary histories of the respective organisms. We suggest that the similar developmental trajectories of Myxobacteria and Dictyostelia are more plausibly due to shared generic physical processes in coordination with analogous agent-type behaviors than to convergent evolution under parallel selection regimes. Finally, we discuss the broader implications of the existence and synergy of these two categories of developmental factors for evolutionary theory.

## INTRODUCTION

The emergence of multicellular organisms exhibiting cell differentiation, spatial patterning and morphogenesis has been recognized as one of the major transitions in evolution (Maynard Smith and Szathmáry, 1995). Depending on the criteria applied (cell–cell attachment, cell communication, division of cell labor, among others) multicellularity evolved on anywhere between 10 and 25 independent occasions (Niklas and Newman, 2013; Niklas and Newman, 2019). The appearance of multicellular organisms enabled an extraordinary increase in the complexity of living systems and the study of the developmental mechanisms and selective forces leading to its emergence, maintenance, and variation is an active research area (e.g., Niklas and Newman (2016). In broad terms, multicellular organisms can be classified either as aggregative (“coming together”) or zygotic (“staying together”), according to the mechanism by which multicellularity arises (Bonner, 1993; Tarnita et al., 2013). In the former, multicellular organisms develop through the gathering of several individual cells potentially belonging to different genetic lineages; in the latter, all the cells in the organism are the offspring of a single cell and remain attached to each other after cell division (Bonner, 1998; Grosberg and Strathmann, 2007). Across eukaryote lineages, aggregative multicellularity involves amoeboid cells and leads to the formation of a fruiting body or “sorocarp” (Brown and Silberman, 2013). There appear to be ecological determinants (e.g., resource availability, land vs. water environment) of whether organisms are clonal or aggregative (Bonner, 1998; Fisher et al., 2019; Hamant et al., 2019). Furthermore, clonal lineages do not always exhibit complex development with different cell types and arrangements, and aggregative ones often do (Newman, 2014b; Newman, 2019c; Niklas and Newman, 2019).

Dictyostelia and Myxobacteria are eukaryotic and prokaryotic multicellular lineages, respectively (Romeralo et al., 2013a; Yang and Higgs, 2014). In these lineages, the life cycle comprises a vegetative and a developmental stage. In the vegetative stage, Dictyostelia behave as solitary cells acting independently of each other, and with the possible exception of intercellular repulsion during feeding (Keating and Bonner, 1977), only engage in cell-cell interactions during development. In contrast, Myxobacteria, often referred to as social bacteria, are believed to organize into cell consortiums through their entire life cycles, although single-cell-specific behaviors are observed in the laboratory (Thutupalli et al. (2015) and unpublished observations). Both lineages are commonly found in soils where they feed upon (other) bacterial species. Once nutrients have been depleted, they transit into a developmental stage characterized by a substratum-dependent cellular aggregation that culminates in the formation of multicellular structures called fruiting bodies, containing up to 10^5^-10^6^ cells, where cell differentiation takes place (Whitworth, 2008).

The basis of cell differentiation in *D. discoideum* has been explained in two ways. There are pre-aggregation tendencies among amoebae, stochastic in origin, biased by the environment they experienced during the phases of growth and division, or, cell differentiation is a post-aggregation phenomenon based on intercellular interactions and diffusible morphogens (reviewed in (Nanjundiah and Saran 1992). There is experimental evidence for each of the two viewpoints (Kawli and Kaushik, 2001), and it is also clear that subsequent interactions can override cell-autonomous tendencies (Raper, 1940).

In Myxobacteria development, cells commit to at least three different cell types, peripheral rods, spores and autolysis. In Dictyostelia, there are principally only two terminal cell types, stalk and spore cells, with several transitory cell types (different pre-stalk and pre-spore subtypes) observed over the normal course of development. Phylogenetic analyses suggest that the capacity for cellular differentiation predated the emergence of multicellular development in both lineages (Arias Del Angel et al., 2017; Schaap et al., 2006). Theoretical studies show that cellular differentiation can spontaneously arise by the coupling of multistable cellular systems (Furusawa and Kaneko, 2002; Mora Van Cauwelaert et al., 2015).

The morphology of fruiting bodies in both lineages displays a similar extent of diversity ranging from simple mound-like to highly branched tree-like structures. Morphology is a species-dependent trait, though there are examples in the dictyostelids of the fruiting body of one species mimicking the morphology of another (Bonner, 2009). For neither Myxobacteria nor Dictyostelia are fruiting bodies morphologies a monophyletic trait (Arias Del Angel et al., 2017; Schaap et al., 2006), and thus different forms are likely to have evolved multiple times within each lineage.

The issue of convergence becomes even more remarkable when it is recognized that sorocarpic amoebae like those of Dictyostelia occur in five of the seven supergroups into which eukaryotes are divided. (Archaeplastids, the group containing red algae, green algae, and plants, appear to be the sole exception. In another supergroup, the Alveolates, aggregative multicellularity and fruiting body formation occurs, but in ciliates, not amoebae (Bonner, 2009; Brown and Silberman, 2013).

Perhaps more surprising is the resemblance of developmental processes and resulting morphologies between eukaryotic sorocarpic amoebae such as Dictyostelia and the prokaryotic Myxobacteria, despite their independent origins, the evolutionary distance between them, and the lack of traceable homology in the molecular mechanisms in each group (Fig. 1). Bonner (1982) suggested that the parallelisms between Myxobacteria and Dictyostelids appear as a consequence of either similar selective pressures or shared developmental constraints. But these determinants are not mutually exclusive and discrimination between them is not trivial (Olson, 2012). Kaiser (1986) proposed that a joint investigation of Myxobacteria and Dictyostelia could potentially lead to the identification of generalities underlying the multicellular phenotypes across both lineages.

**Figure 1.**
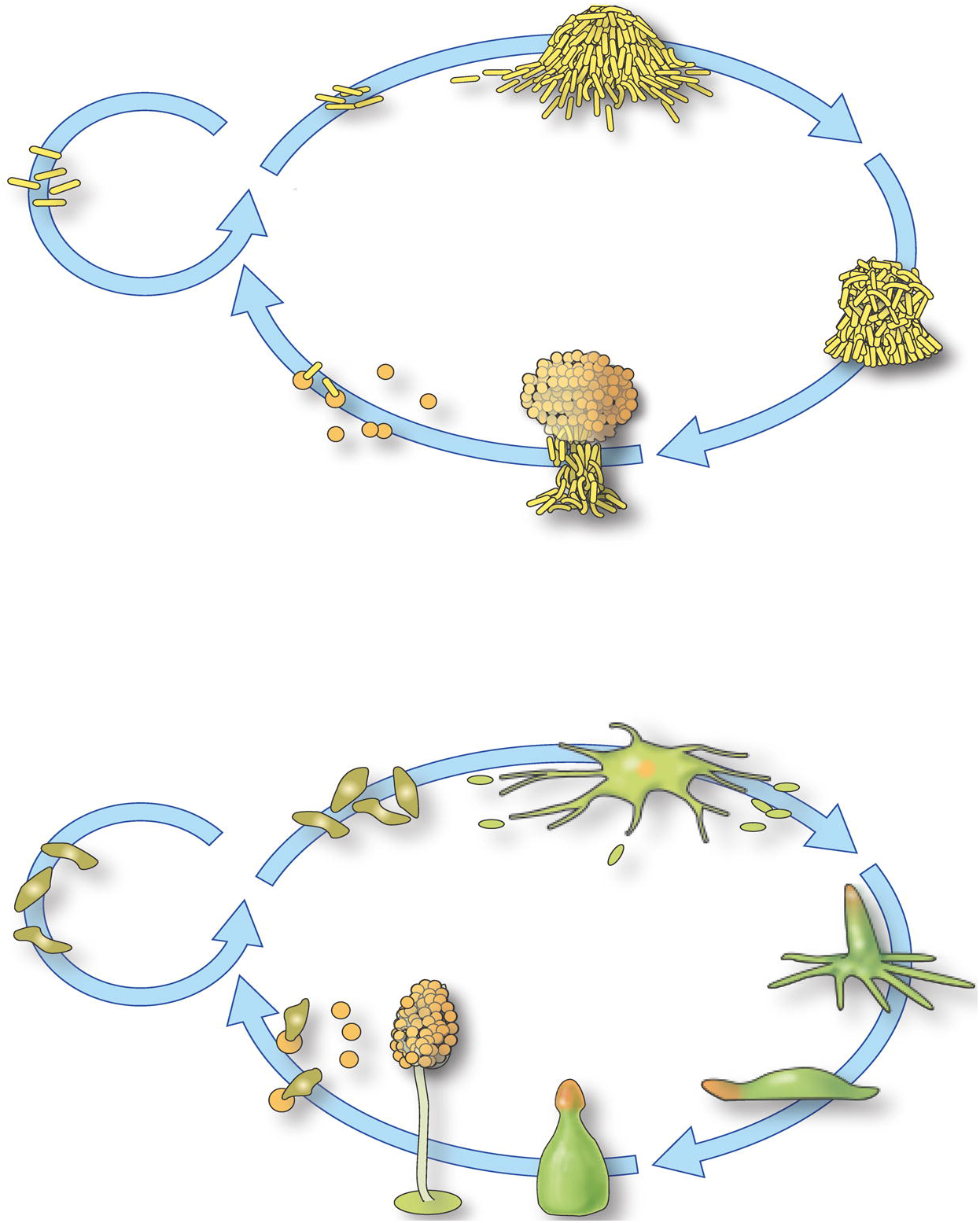
(Upper panel) Life cycle of *Myxobacteria xanthus*, a representative multicellular myxobacterium. The circle on the left represents the proliferative mode that occurs in a nutrient-replete setting. The oval on the right shows the sequence of stages initiated under conditions of starvation: clockwise, from top left, aggregation, mound formation, fruiting body formation and spore differentiation. Spores can be dispersed and may germinate as single vegetative cells under nutrient-rich conditions. (Lower panel) Life cycle of *Dictyostelium discoideum*, a representative dictyostelid. The circle on the left represents the proliferative mode that occurs in a nutrient-replete setting. The oval on the right shows the sequence of stages initiated under conditions of starvation (clockwise, from top left: starved amoebae, developing aggregation, late aggregations, migrating slug, developing fruiting body, finished fruiting body with spore mass supported by an erect stalk, amoebae emerging from spores after dispersal).

Since Kaiser’s proposal, a combination of experimental and modeling approaches has been employed to investigate the development in these two lineages (Romeralo et al., 2013b; Yang and Higgs, 2014). Such studies advanced after physico-chemical processes came to be considered as key factors determining the developmental outcomes (Bretschneider et al., 2016; Fujimori et al., 2019; Thutupalli et al., 2015; Umeda and Inouye, 2002). Specifically, there is a recognition that the shaping of multicellular masses cannot be explained independently of their material properties, and that developing organisms are thus subject to physical forces and effects relevant to their composition and scale (Benítez et al., 2018; Newman, 2014a; Newman and Bhat, 2009; Rivera-Yoshida et al., 2018). When applied, for example, to embryonic animal tissues, which behave similarly, in certain respects, to non-living liquids and liquid crystals, physical models predict the formation of immiscible layers, interior spaces, and, when the subunits are anisotropic, the capacity to undergo elongation (Newman and Bhat, 2009). In contrast, plant tissues, characterized by rigid cell walls, behave like deformable, mechanically and chemically active solids which (unlike liquid-state materials) can bud or branch (Benítez et al., 2018).

Properties shared by cellular masses with (as the case may be) nonliving liquids, solids, or semisolid materials have been termed “generic” (Newman and Comper, 1990), and we adopt that term here. The physical forces, effects and processes inherent to such materials enable and constrain developmental outcomes in multicellular masses, leading to the conclusion that homoplasy (the same form, independently evolved) is expected to be common, and some morphological motifs should be recurrent and predictable (Benítez et al., 2018; Newman, 2014a). Physical determinants, in this view, are complementary to the regulatory dynamics within cells. Indeed, physical and physicochemical processes are mobilized on the multicellular scale by genes, their products and other molecules, and are thus subject to regulation throughout evolution (Benítez et al., 2018).

In contrast to the molecular subunits of non-living materials, the individual cells constituting a multicellular cluster are able to sense and respond to local cues through signaling and regulatory pathways. Because of their intracellular chemical dynamics and capacity to generate mechanical forces, cells can be understood as agents that actively modify their behavior in response to their environment, and even modify their environment in ways that can further affect the cell-environment interaction. These processes taking place at the cell level, including chemotaxis, which as discussed in Section 4, can continue even when the cells are already aggregated, can translate into collective behaviors that act in parallel and coordination with, and even oppose, the generic physical processes that shape a tissue mass. These “agent-like” behaviors modify the outcomes that would be expected if only generic physical processes were operative.

Here, we hypothesize that the morphological outcomes, and thus the parallelism between the myxobacterial and dictyostelid lineages, originated as a consequence of the interplay between generic processes acting upon the multicellular materials and agent-like behaviors characteristic of the constituent cells. To this end, we describe the major generic and agent-like properties exhibited during the development of these lineages and attempt to analyze their contributions to the emergence of the groups’ shared traits. We suggest that as a consequence of aggregation, the nascent multicellular mass becomes subject to new sets of patterning and morphogenetic processes resulting from the fact that cell-cell contacts or immersion in a matrix mediate the emergence of a fluid-like properties. In both lineages, this leads to developmental processes, e.g., streaming, rippling, that are similar to behaviors observed in non-living fluids. We explore the idea that deviations of the dynamics and morphological outcomes of the multicellular mass from the generic predictions are due to the contribution of agent-like behaviors of individual cells, e.g., gradient sensing, directed migration, quiescence.

Generic effects are *common causes* in the different lineages. This is because whatever molecules underlie the realization of properties such as cell-cell adhesion, spatial heterogeneity via diffusion gradients, and so in in different lineages, the morphological outcomes are similar by virtue of being produced by similar physical generative processes. Agent-behaviors, in contrast, are peculiar to disparate lineages (cell locomotion, for example, has very different physical and genetic bases in prokaryotes and eukaryotes, as does entry into the quiescent state), reflecting the evolutionary histories of the respective organisms. However, these behaviors can be analogous to one another, thus contributing to convergent morphological outcomes. Further, analogous intracellular dynamical behaviors such as biochemical oscillation can be organized by generic effects such as synchronization, leading to additional shared generic modes of organization. We conclude that the similar developmental programs of Myxobacteria and Dictyostelids are plausibly due to shared generic physical processes in coordination with analogous agent-like behaviors.

## GENERIC MATERIAL PROPERTIES OF MYXOBACTERIAL AND DICTYOSTELID MULTICELLULAR MASSES

Based on the observation that animal life is characterized by a restricted set of basic forms and patterns, Newman and co-workers advanced the conceptual framework of “dynamical patterning modules” (DPMs). DPMs are defined as sets of gene products and other molecules in conjunction with the physical and physicochemical morphogenetic and patterning processes they mobilize in the context of multicellularity (Newman and Bhat, 2008; Newman and Bhat, 2009). These include phenomena such as adhesion and differential adhesion, and reaction-diffusion effects. This framework emphasizes that the material nature of a developing organism makes it subject to generic physical processes (i.e., those common to living and nonliving viscoelastic and excitable systems) and that they readily exhibit morphological motifs – layers, segments, protrusions – Inherent to the respective materials. The term “module” is employed to highlight the semi-autonomous action of DPMs in determining specific spatial patterns and structures. But the DPMs also interact during development and can thus be conceptualized as a complex “pattern language” for generating organismal form (Hernández-Hernández et al., 2012; Newman and Bhat, 2009). This approach is distinguished from a purely “tissue physics” framework since it also recognizes the relevance of the cells as repositories of genetic information, making such systems subject to evolutionary processes not applicable to non-living matter.

Even when the similarity in the mesoscopic (i.e., physics of the middle scale) properties of living and certain kinds of non-living matter is recognized, it should not be taken to imply that they are constituted in the same way. The liquid or solid nature of living tissues does not arise from the same subunit-subunit interactions that endow non-living materials with these properties. This is particularly the case with the liquid-like state of animal tissues. Instead of the thermal vibration-driven Brownian motion that causes the molecular subunits of non-living liquids to move randomly, the cells in animal tissues move actively by ATP-dependent cytoskeleton-generated forces, which in the absence of external signals is also random. Despite continually changing their neighbors, subunits of nonliving liquids cohere due to the weakly attractive electronic interactions that hold them together. The cells of developing animal tissues also remain cohesive despite their translocation, but for a different reason: the homophilic attachment proteins (classical cadherins) that mediate their transient attachment extend through the cells’ membranes to form stable connections between adhesive and motile functions (Newman, 2019b). In plant and fungal tissues, instead of the charge-based or covalent bonds of the atomic or molecular subunits of non-biological solids, the cells are cemented together by pectins and glycoproteins which are subject to unique forms of reversible remodeling (Benítez et al., 2018; Hernández-Hernández et al., 2012). Because these generic properties are dependent on evolved biological, rather than purely physical effects, the various viscoelastic and deformable solid materials that constitute living tissues have been termed “biogeneric” matter (Newman, 2016).

In the following, we describe some of the generic and biogeneric properties and processes of Myxobacteria and Dictyostelia multicellular masses and compare these properties to those implicated in animal development. Then, we describe the molecular components that establish and mobilize these properties in both Myxobacteria and Dictyostelia. Next, we highlight some developmental phenomena in these organisms and evaluate the extent to which these can be explained by generic physical behaviors, and what is left unaccounted for. Finally, we describe the agent-like behaviors of the subunits (bacteria and amoeba) of the two systems, discuss their similarities and differences, and discuss how analogous agent behaviors coordinate with and complement the described generic properties, and potentially account for the common developmental modes of Myxobacteria and Dictyostelia.

### Adhesion- and matrix-based cell-cell association

Cell adhesion is the defining characteristic of multicellular organisms and the nature and strength of cell bonding is a major determinant of tissue properties (Forgacs and Newman, 2005; Mora Van Cauwelaert et al., 2015; Niklas and Newman, 2019). In animals, cell-cell adhesion is mediated by membrane proteins such as cadherins that permit cells to be independently mobile and capable of moving relative to another while remaining cohesive. As noted above, the animal tissues from which embryos and organs develop behave formally like liquids (Newman, 2016).

In *D. discoideum*, cell-cell adhesion at early stages of development involves the action of several proteins including the immunoglobulin-like DdCAD-1 and the glycoproteins gp80 and gp150 whose expression and activities are tightly regulated during the different stages of development (Coates and Harwood, 2001). Later in development, when cells have entered into streams and cell density has increased, the cells are also embedded in cellulose-based matrices that provide the basis for adhesion in cellular conglomerates (Huber and O’Day, 2017). In the case of *M. xanthus*, persistent cohesion is correlated with the secretion of thick fibrils, composed of carbohydrates and proteins, that coat the cell surface and constitute an extracellular matrix that interconnects the cells (Arnold and Shimkets, 1988; Behmlander and Dworkin, 1994a; Behmlander and Dworkin, 1994b). Chemical or genetic disruption of fibrils causes defects in agglutination and failures in social and developmental behaviors. In a similar fashion to the animals, cell-cell adhesion in Myxobacteria and Dictyostelia depend, to different degrees, on the presence of divalent cations (Lin et al., 2006; Shimkets, 1986).

Both Myxobacteria and Dictyostelia also have strong associations with external substrata during their pre-culmination stages of development. The closest analogy in animal systems is the interaction of cell layers in eumetazoans with internally generated planar basal laminae, which are not generally present in the earliest diverging and morphologically simplest metazoans, sponges and placozoans. In both Myxobacteria and Dictyostelia cells are more loosely associated with one another as they interact with the substratum than are the cells in planar animal epithelia. In the non-animal systems, cell substratum interactions depend on focal adhesions that indirectly (in contrast to directly in animal tissues) mediate communication between the substratum and the actin cytoskeleton, where they also provide the foundation for cellular motility (Faure et al., 2016; Fukujin et al., 2016).

A key difference between the respective lineages is that dictyostelid cells only engage in persistent cell-cell interactions shortly after starvation, whereas extensive cell-cell adhesion and interactions take place among myxobacterial cells through their entire life cycle. While the mechanisms involved in cell-cell and cell-substratum contact between Myxobacteria and Dictyostelia are different, in both cases the bonds between adjacent cells are weak enough to allow cells to rearrange relative to one another during aggregation and shortly after mounds are formed. Therefore, aggregating cells in these lineages behave like non-living liquids, exhibiting streaming and rippling behaviors characteristic of such materials. This contrasts with monolayered animal tissues (epithelia) which, though also having liquid-like properties in the plane, bind too strongly to their intra-organismal substrata, *basal laminae*, to manifest similar fluid-like behaviors at the planar interface (Mittenthal and Mazo, 1983).

Unlike Dictyostelia, in Myxobacteria some type of cell-cell adhesion or matrix embedment is present throughout the whole life cycle, causing cellular masses to exhibit liquid-like behaviors in both vegetative and developmental stages (Thutupalli et al., 2015). During predation, cells align and move concertedly into ripple-like travelling waves (Zhang et al., 2012). Once development has started, *M. xanthus* aggregation is largely driven by entropy minimization through reduction of the surface area on which the collective cell population contacts the substratum (Bahar et al., 2014). This is a comparable behavior to that of liquid droplets, where individual subunits or clusters move into larger droplets of larger volume but smaller contact area with the surface. In Myxobacteria, phase separation has not been implicated in sorting of cell types inside fruiting bodies. However, since spores are coated by material that increases cell cohesiveness, differential adhesion likely contributes to the spontaneous sorting out of spores from peripheral rod cells, reflecting their liquid-like properties.

It is important to distinguish the liquid-like properties of both Dictyostelia and Myxobacteria cell streams and masses from that of embryonic animal tissues. In epithelioid animal tissues the cells are directly attached to their neighbors by transmembrane cadherins which maintain strong cohesivity while permitting rearrangement. This is consistent with persistent apicobasal polarization that allows for the formation of lumens within cell masses, and planar cell polarization that permits elongation and other reshaping of tissues by intercalation and convergent extension, a liquid-crystalline like phase transformation. In Dictyostelia, the cells are embedded in cellulose-based matrices that enable cell rearrangement and hence the liquid-like behaviors described above (Huber and O’Day, 2017). However, the lack of direct engagement in this attachment mode with the cytoskeleton makes cell polarization, even when it occurs, transient and unconducive to lumen formation or stable intercalation (Manahan et al., 2004; however, see Hayakawa et al. (2020)). Cells of Dictyostelia also have a more pronounced chemotactic response to extracellular signals than most animal embryonic cells, which contributes to their particular version of liquid-tissue properties (Tan and Chiam, 2014) (see below).

The glycoprotein-based associations of Myxobacterial cells are also too transient, and their polarity too rapidly reversible, to allow lumens to form, at least until solidification occurs during fruiting body formation (see below). However, the cells are stably elongated by default, and thus readily form liquid crystalline-like domains as in some animal tissues (Thutupalli et al., 2015). The rapid relative movement of the cells, though, causes these to be only local and temporary.

### Solidification

The generic-type fluid-to-solid transitions seen during development of the aggregative species can productively be considered in relation to well-studied ones in animal embryogenesis. Animal tissues during early stages of development, as noted above, behave in important ways like non-living liquids. As development proceeds, however, some tissues undergo a transformation where cell movements become constrained and the cellular mass behaves more like a solid (Newman, 2019b). In these tissues, solidification may provide increased mechanical integrity, and new morphological outcomes and constructional elements (e.g., exo- and endoskeletons) arise with the physical properties of these materials. The most typical way solidification occurs is by the deposition of stiff extracellular matrices (ECM), consisting of fibrous and nonfibrous proteins such as collagen and elastin, covalently linked to, or complexed with glycosaminoglycan-type polysaccharides. These ECMs can also become mineralized, as in bone and tooth. More recently, “jamming,” a liquid-to-solid transition known from colloid physics (Bi et al., 2011) has been shown to occur in liquid-state tissues as a result of increased cell-cell adhesivity (Mongera et al., 2018).

In *D. discoideum*, cells are embedded in an ECM that once aggregation is complete defines the boundaries of the aggregate. Aggregation in this and related species leads to the formation of a migratory “slug” (see below), which once it reaches its final position, forms a fruiting body by building up a stalk that takes cellular material away from the surface, and in which terminal cell differentiation takes place. Membrane proteins involved in cell-cell adhesion are expressed in a cell-type dependent fashion. Spores and stalk cells phase separate, in part, due to the resulting differential adhesion, in agreement with the expected behavior of immiscible liquids (e.g., water-oil mixtures), although other factors such as chemotaxis and differential cell motility are also involved (see below) (Bretschneider et al., 2016; Raper, 1940).

During fruiting body elevation deposition of ECM, is required for the stiffening and construction of the stalk (Palsson, 2008) (Dickinson et al., 2012). Solidification occurs unevenly across the cellular mass. While the movement of cells in the stalk becomes constrained because of the ECM, the remaining cells move upwards as the stalk continues to be built up following the expected dynamics of solidifying non-living liquids. In Myxobacteria, deposition of a stiff ECM appears to be the most important factor in aggregation, but the “solidification” of maturing fruiting bodies may also involve jamming (Hu et al., 2012; Liu et al., 2019; Thutupalli et al., 2015) see below).

### Differential loss of mass

In animal morphogenesis, differential loss of mass can be achieved through programmed cell death (e.g., apoptosis, autophagy and necrosis) where, in addition to acting as cue for signaling pathways, it can also induce tissue reshaping by cell elimination or mobilization of mechanical forces (Monier and Suzanne, 2015; Suzanne and Steller, 2013). In both Myxobacteria and Dictyostelia, it has been suggested that programed cell death may act as a mechanism for nutrient release and recycling that can be employed for the remaining cells in the population as source of energy and cellular materials (Boynton et al., 2013; Mesquita et al., 2017). However, localized developmental lysis may also be relevant in mechanical reshaping multicellular microbial masses. For example, localized cell death mobilizes mechanical forces that are instructive for the generation of key features during development of *B. subtilis* biofilms (Asally et al., 2012). In Myxobacteria, where most of the cells in the initial population undergo developmental lysis, lysed cells may serve to strengthen the ECM (Hu et al., 2012). Specifically, exopolysaccharides embedded in the ECM interact with extracellular DNA. As a consequence, the ECM exhibits greater strength and stress resistance (Hu et al., 2012). While the origin of this extracellular DNA remains unclear, it may be released by cells after lysis. In Myxobacteria and Dictyostelia, peripheral rods and stalk cells, respectively, die after the stalk has been built up. In both, cell death is a consequence of nutrition deprivation. In the dictyostelids, it shows similarities as well as differences with the manner in which cell death is regulated in metazoan tissues (Arnoult et al., 2001; Cornillon et al., 1994; de Chastellier and Ryter, 1977; Kawli et al., 2002).

Additional generic effects can arise in cell masses from, e.g., synchronization of intracellular biochemical oscillations. Some of these will be characterized below, after the roles of such pivotal cellular functions in individual cell behavior are described.

## AGENT-LIKE BEHAVIORS IN MYXOBACTERIA AND DICTYOSTELIA

Previous descriptions of the development of embryonic animal and plant tissues in terms of material properties of multicellular assemblages have accounted for key morphological features on the basis generic physical processes pertaining to these materials without invoking the idea that individual cellular subunits of such materials act as autonomous agents in creating multicellular forms and patterns (see, e.g., (Benítez et al., 2018; Newman, 2016). Although the constituent cells in these generic accounts are assumed to carry out metabolic and synthetic functions necessary to sustain life, to change their state (including polarity) in response to external signals (Niklas et al., 2019), and (in the case of animal systems) locomote randomly, the materials-based perspective does not involve formal sets of rules governing cellular interactions of individually mobile cells. Similarly, as seen in the previous section, several important aspects of Myxobacteria and Dictyostelia development can be explained by considering them as generic materials, that is, considering the cell streams and masses as generic liquid-like or solid-like materials.

However, attempts to computationally model aggregation of Myxobacteria and Dictyostelia cells and the resulting multicellular masses based on generic mesoscale physics have found the need to incorporate agent-like behaviors of the cells themselves into the models to capture the relevant behaviors (Bahar et al., 2014; Fujimori et al., 2019; Marée and Hogeweg, 2001; Thutupalli et al., 2015). (Following standard usage (Thorne et al., 2007) we define agents as autonomous entities acting according to internal rules in a shared environment.) For biological agents such as Myxobacteria and Dictyostelia cells these “rules” depend on intracellular dynamics of molecules and pathways.

In biological development, agent-based phenomena pertain to the semi-autonomous activities of individual cells or cells in transient associations with each other. This contrasts with the collective effects governed by generic physical processes operating at the mesoscale. Unlike nonliving systems, the subunits of tissues, aggregates, and presumptive aggregates are living cells that are internally complex and chemically, mechanically, and electrically active and potentially excitable. Cell dynamics can modulate the properties of biomaterials, making a liquid-like animal tissue liquid-crystalline, for example, or a solid plant tissue locally expansible. When cells act as individuals, however, alterations in their internal states can give them agent-like properties when interacting with other such agents or features of the environment. The reality of this distinction is illustrated by a recent study of neural crest migration where, exceptionally in animal systems, cells navigate directionally through surrounding tissues in loose association with each other. Consequently an agent-based modeling approach was deemed necessary (Giniunaite et al., 2020a).

In certain cases, generic properties and agent-like effects mobilize the same intracellular activities and processes. For instance, random cell movement, driven by actomyosin-based contractile and protrusive activity, is essential to the liquid-like state of animal tissues. These processes in individual amoeboid cells can also be mobilized for directional locomotion. Similarly, concerted induction of cell polarity in animals and plants can impart anisotropy to the respective tissues, changing their shapes and topology (Nance, 2014; Niklas et al., 2019). In single amoeboid or bacterial cells, in contrast, polarity is essential in the sensing of chemical and substrate gradients and directed navigation. Lastly, intracellular biochemical oscillation in animal, amoebal, or bacterial cell collectives can attain synchrony, thereby causing it to behave as a “morphogenetic field” in which cell states are coordinated at long distances across the multicellular mass (Bhat et al., 2019 and references below).

As described above, multicellular systems can exhibit predictably similar morphological and patterning outcomes as a result of mobilizing generic mesoscale physics. Agent-like behaviors, however, are not generic in the same in sense, and their outcomes do not have the same kind of shared inherency, since the rules that individual cells follow in relating to other cells and their external environments are specific to each lineage and dependent on their respective evolutionary histories. As mentioned above, and exemplified in the phenomena of directed migration, regulated quiescence, and oscillation-based cell-cell communication, agent-like behaviors of cells as distantly related as Dictyostelia and Myxobacteria can sometimes have analogous morphological outcomes. This, combined with the generic effects with which they interact in the development of multicellularity, contribute to the strikingly similar morphological motifs in these disparate systems.

### Directed migration

During animal embryogenesis, the displacements of cells relative to another can be largely understood in terms of random movements analogous to the Brownian motion of the molecular subunits of non-living liquid systems (Newman and Bhat, 2009). In Dictyostelia and Myxobacteria, in contrast, cell trajectories deviate from the undirected motion of most animal tissues due to the action of signaling and regulatory mechanisms. These bias the direction and speed of cell movement in response to local cues in ways that may change as development progresses. We suggest that some particularities of Dictyostelia and Myxobacteria observed at the mesoscale (notwithstanding their shared liquid-like behaviors) derive from the distinct mechanisms underlying directed cell migration in these two groups.

In Dictyostelia, cell movement occurs by amoeboid motion, which is driven by cytoplasmic actomyosin-based contractile and protrusive activity just as in animal cells (Fukui, 2002). In contrast to the generally random cell locomotion seen in animal tissues, however, Dictyostelia exhibit both random movement and directed movement via chemotaxis, which can be thought of as a biased random walk. Amoebae seek food by chemotaxis. Aggregation is also mediated by chemotaxis, but to an aggregation pheromone (e.g., cAMP). Chemotaxis remains essential for all subsequent developmental stages (Du et al., 2015). It dependent on both the physical process of diffusion of the chemoattractant (which is not a generic tissue mechanism since it is outside the cell mass) and agent-like behavior in response to the chemoattractant signaling at the cellular level. Specifically, chemotaxis is a quantifiable outcome of directional pseudopod extension (Chopra and Nanjundiah, 2013).

In *D. discoideum*, the response to the chemoattractant cyclic AMP (cAMP) involves an oscillatory dynamics of excitation and adaptation (see below). The formation of streams with high cellular density is facilitated by the collective movement of cells coordinated by chemotaxis towards higher concentrations of cAMP. While cellular movements are most prominent at the aggregation stages, extensive cell translocation still take place at later stages of the development with chemotaxis biasing the individual movements. Cell movements remain operational in the concerted movement of cells within a slug (Singer et al. (2019) but see Hashimura et al. (2019). Finally, in slugs and maturing fruiting bodies, chemotaxis operates jointly with differential adhesion to drive cell sorting (an authentically generic tissue process) where it also provides the basis for fruiting body elongation (Matsukuma and Durston, 1979; Schaap, 2011; Tan and Chiam, 2014).

In the case of Myxobacteria, where cells are rod-shaped, the presence of protein complexes that promote motility defines a lagging and a leading pole (Guzzo et al., 2018). Cells in transient contact with their neighbors move along their long axis in the direction of the leading pole, with reversals in the direction of movement being a major agent-type behavior in Myxobacteria motility. Reversals occur by switching the cellular polarity (i.e., the leading pole turns into the lagging pole and vice-versa) and net cellular displacement is influenced by the reversal frequency (Cotter et al., 2017). At the molecular level, reversals are controlled by the Frz and MglAB intracellular oscillators (Guzzo et al., 2018; Igoshin et al., 2004). Directed migration is favored during development by a reduction in the frequency of reversal that allows cells to retain their direction and aggregate. This frequency reduction is stimulated by cell-cell contacts, likely involving the exchange of intercellular signals, which become more frequent as aggregation proceeds and cellular density increases (Cotter et al., 2017; Zhang et al., 2018). An additional mechanism underlying directed migration in Myxobacteria is *stigmergy*, by which individual cellular movement is biased by cues left behind by other cells (Gloag et al., 2016). Specifically, while moving over solid surfaces, *M. xanthus* cells deposit slime material that forms trails over which other cells travel preferentially.

In both Myxobacteria and Dictyostelia, the interplay between directed migration, an agent-like behavior, and generic material properties highlights the need to consider them together in accounting for development. In *D. discoideum*, cell sorting requires agent-like behaviors (directed migration) and generic properties (differential adhesion) for its completion. In Myxobacteria mesoscopic movement patterns are the result of the joint effect of the agent-like behavior of directed migration and generic liquid-like behavior enabled by transient cell-cell adhesion. In addition to these, the different phenomena observed along Myxobacteria life cycle also require cellular alignment that may occur spontaneously as a generic property of rod-shaped particles and cells (Janulevicius et al., 2015; Volfson et al., 2008).

### Cessation of movement and quiescence

Development in *M. xanthus* and other myxobacteria starts as a response to starvation (Dworkin, 2007). Once it is sensed, ribosomes stall and the enzyme RelA increases the intracellular concentration of the tetra- and pentaphosphate alarmones (p)ppGpp which, as in most bacteria, induces the so-called stringent response (SR; (Boutte and Crosson, 2013; Cabello et al., 2017; Chatterji and Ojha, 2001; Manoil and Kaiser, 1980a; Manoil and Kaiser, 1980b; Shimkets, 1999). As (p)ppGpp accumulates, proteases are synthesized and exported, leading to an extracellular mixture of amino acids and peptides (A-signal), where it mediates a quorum-sensing mechanism that enables a coordinated population-level response to starvation, including specifying the minimal cell density required for initiation of development (Kuspa et al., 1992). Myxobacteria respond to nutrient depletion via the SR, but also require high cell density to initiate fruiting body and spore development. To effect this, in addition to conserved SR components, Myxobacteria produce CgsA, which positively regulates (p)ppGpp and is in turn positively regulated by it, and SocE, which suppresses and is suppressed by the production of (p)ppGpp (Boutte and Crosson, 2013; Crawford and Shimkets, 2000a; Crawford and Shimkets, 2000b). Therefore, when A-signal rises to the concentration where it promotes aggregation (Bretl and Kirby, 2016), which in non-aggregative bacteria would turn off the SR (since the A-signal components serve as nutrients), the downregulation of SocE permits CgsA to keep (p)ppGpp (which is required for spore formation) elevated during development.

A proteolytic cleavage product of CsgA serves as another extracellular signal which is required for fruiting body development and sporulation (C-signal; (Giglio et al., 2015; Gronewold and Kaiser, 2002). The specific mechanisms by which C-signal mediates intercellular communication are not understood, but it appears to be involved in cell-to-cell adhesion and coordination of cell movement during development (Sogaard-Andersen et al., 2003) and is a key element enabling multicellular aggregation and cellular differentiation (Holmes et al., 2010; Julien et al., 2000). In addition to A- and C-signaling, at least three other signals, termed B-, D- and E-signal, mediate intercellular communication and coordination of individual cells during development, but their specific mechanisms remain unclear (Bretl and Kirby, 2016; Kaiser, 2004).

The SR is largely conserved in bacteria where it typically mediates proliferative and biosynthetic quiescence in response to nutrient depletion and other stresses. While it is therefore likely to have been present in the unicellular ancestor of myxobacteria, the genetic novelties represented by the intracellular CsgA-SocE circuits and the extracellular A-, B-, C-, D- and E-signals co-opted this behavior to the transition to multicellularity. By making the SR cell nonautonomous, these components and their interactions form a set of rules that enable cells of *M. xanthus* to act as agents with respect to both cessation of movement and active signaling (Arias Del Angel et al., 2017). As demonstrated in other myxobacteria such as *Anaeromyxobacter dehalogenans*, and *Sorangium cellulosum*, it likely maintains aggregates and promotes the differentiation of their constituent cells into quiescent spores and other cell types (Huntley et al., 2014; Knauber et al., 2008).

Eukaryotic cells like those of Dictyostelium do not have a bacterial-type stringent response, but they have their own conserved sensor of nutrient depletion, the enzyme AMP-dependent protein kinase (AMPK). Among other effects, AMPK inhibits the energy utilization hub mechanistic target of rapamycin complex-1 (mTORC1) under starvation conditions (Hardie, 2014). In animal systems AMPK plays developmental roles in, for example, inducing quiescence in germline stem cells (GSCs) in the nematode *Caenorhabditis elegans*. In the absence of AMPK, the GSCs overproliferate and lose their reproductive capacity, leading to sterility (Kadekar and Roy, 2019). Significantly, in relation to the discussion above of the SR in Myxobacteria quiescence, the function of AMPK in *C. elegans* development has been reconfigured evolutionarily to be cell nonautonomous, with AMPK activity in somatic cells being transmitted to GCSs via small RNAs (Kadekar and Roy, 2019). But the quiescence-inducing role of AMPK is conserved across the eukaryotes, also appearing in plants and fungi (Guerinier et al., 2013; Zhang and Cao, 2017).

In Dictyostelia, AMPK was found to regulate aggregate size and patterning, as well as cell fate choice and stalk-spore case boundary formation in the fruiting body (Maurya et al., 2017). Deletion of the gene specifying AMPK resulted in generation of numerous small-sized aggregates (compared to wild type cell populations) that develop asynchronously to form few fruiting bodies with small spore masses and long stalks. In contrast, when the gene is overexpressed, cells form fruiting bodies with small stalks and large spore masses (Maurya et al., 2017). Although AMPK itself functions cell autonomously, its regulation depends on interaction with other cells, mediated by soluble factors. For example, the secreted inhibitor of cell-cell adhesion Countin (Jang and Gomer, 2008) is upregulated in AMPK null cells, and conditioned media collected from them cause wild-type cells to form smaller aggregates (Maurya et al., 2017).

As with Myxobacteria, the starvation response triggers development at the expense of growth. Jaiswal and coworkers have shown that although in Dictyostelium, mTORC1 function is indeed inactivated via AMPK upon starvation, development is nonetheless initiated. These investigators have identified of a class of essential starvation-upregulated, developmentally associated signaling genes and downregulated growth genes (Jaiswal and Kimmel, 2019; Jaiswal et al., 2019). Based on the earlier work of Maurya et al. (2017), downregulation of the paracrine adhesion inhibitor Countin appears to be a component of this response, suggesting as with Myxobacteria, a conserved starvation-sensing mechanism may have been recruited into a mechanism of multicellular development by one or more factors that mediate communication among agent-like cells.

## OSCILLATIONS AS A BASIS FOR BOTH GENERIC AND AGENT-TYPE BEHAVIORS

Both Myxobacteria and Dictyostelia exhibit intracellular oscillations, which in the first case mainly involves cell polarity and direction of motion reversals, and in the second, production of chemoattractant molecules such as cAMP. Oscillations can mediate global effects if they come into synchrony in established cell masses. This produces developmental fields in which the constituent cells acquire a uniform state in a key modulator (e.g., the transcriptional coregulator Hes1) and therefore are poised to respond to developmental signals in a coordinated fashion. This occurs in animal systems, for example during the formation of somites, tandem blocks of tissue along the central axis of vertebrates (Hubaud and Pourquié, 2014), and the digits of the tetrapod limb (Bhat et al., 2019). The synchronization of oscillators can be considered a generic physical effect since its physical basis is the same regardless of the underlying basis of the oscillation.

But oscillations of individual cells can also provide component of agent-like behavior, particularly in species that develop by aggregation. For example, they can permit cells to signal one another over distances provided they are specifically receptive to periodic stimulation. The myxobacterium *M. xanthus* exhibits a quasi-periodic reversal in the direction of motion. Reversal in the gliding cells are achieved by dynamic cell polarity that switches direction by 180° (Zusman et al., 2007). As noted above, regular reversals are driven by the relocalization of polarity and motility proteins between the leading and lagging poles of the cells and allow for diverse collective modes, such as rippling in nutrient-rich media (Mauriello et al., 2010; Shimkets and Kaiser, 1982). Reversals also appear to be critical for complex collective behavior before and during development (Blackhart and Zusman, 1985; Wu et al., 2009).

Indeed, it appears that reversal frequency in *M. xanthus* drives a phase transition from two-dimensional flocking to one-dimensional streaming, therefore modulating the complex behaviors that enable the robust formation of fruiting bodies (Thutupalli et al., 2015). Because the reversal is coupled to intercellular signaling pathways (C-signal), this periodic switch may be synchronized between different cells and favor development (Igoshin et al., 2004). A refractory period, i.e., time lag in response to the environmental signal(s), in the molecular circuit responsible for inducing the polarity reversal, has been proposed to underlie the rippling dynamics of the bacterial sheet (Guzzo et al., 2018).

As in Myxobacteria, oscillations mediate collective behaviors in Dictyostelia, but they are also the basis of agent-like behaviors in these social amoebae. Initially isolated cells of *D. discoideum* aggregate by chemotactic movements in response to the release of periodic pulses of cyclic AMP, which they also amplify and relay. Specifically, when stimulated with extracellular cAMP, cells respond by synthesizing and secreting more cAMP. This results in non-dissipating waves of cAMP which guide aggregation of individual amoeboid cells (Tomchik and Devreotes, 1981). The relay requires a refractory period, or else there could just be an explosive production of cAMP with no local gradients to guide cells into aggregates. So, a nonconstant, ultimately periodic, production of the chemoattractant by the dispersed cells is intrinsic to the patterning process.

Since the cells in this organism start out as individuals, a key question in characterizing their agent-like behavior is the relation of single cell oscillations to the global oscillations in the organizing field of cells (Nanjundiah and Wurster, 1989). Isolated cells are capable of oscillating (Satoh et al., 1985), but it has been unclear whether such oscillations initiate the propagating waves in the “excitable medium” constituted by the field of cells (Cohen and Robertson, 1971; Durston, 1973). There are two physical possibilities. In the first, a set of oscillators (the amoebae in this case) with identical period, but randomly distributed phases come into synchrony or attain a spatiotemporal propagating mode through weak coupling, by a diffusible chemical, for example (Garcia-Ojalvo et al., 2004; Kuramoto, 1984; Strogatz, 2003). The second possibility is that cells only become oscillatory as a result of collective interactions, the global behavior being an emergent process. Gregor et al. (2010) investigated these possibilities experimentally and via mathematical modelling, and while they confirmed that isolated cells are capable of oscillating, they concluded that the second possibility, what they term “dynamical quorum sensing,” was the way that globally synchronized waves are generated in Dictyostelium.

## INTERPLAY OF GENERIC PROPERTIES AND AGENT EFFECTS

As we have shown, aggregative multicellular systems can change their organizational states as a result of the cell masses they form being shaped and reshaped by mesoscopic physical effects, and also by lineage-specific, “custom-built” agent-like behaviors. A schematic representing some of these factors and determinants is shown in Fig. 2. In some cases, however, developmental transformations cannot be attributed to either category of effect alone but can only be understood as outcomes of a combination of the two acting in concert. A newly characterized example of this described by Hayakawa et al. (2020), in which an ordered, liquid-crystalline-like field of polarized *D. discoideum* amoebae organizes by phase separation, from populations of cells of a mutant strain incapable of chemotactic signaling via cAMP. This novel patterning phenomenon, which has generic-type features, occurs by “contact following locomotion,” a behavior whose agent-type role in the collective motion is supported by simulations.

**Figure 2.**
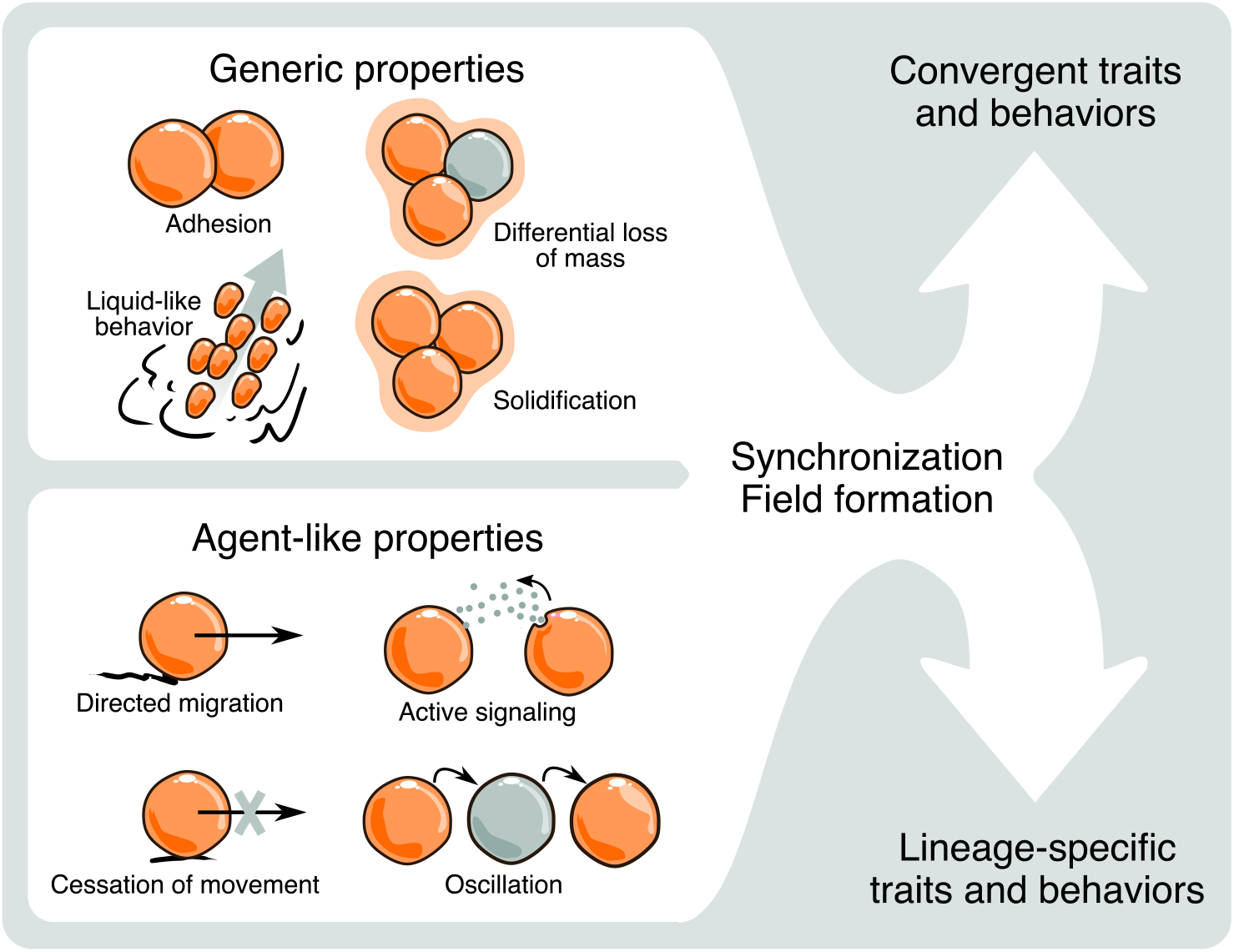
Schematic representation of (left, top) a selection of generic physical effects and one of their underlying mediators (cell-cell adhesion), and (left, bottom) a selection of agent-like effects, all of which pertain to aggregative multicellular organisms such as myxobacteria and dictyostelids. Some individual cell behaviors like biochemical or polarity oscillation can, when they operate in the multicellular context, can mediate global generic effects, like morphogenetic fields in which cell state is coordinated over large distances. Generic processes can lead to convergent morphologies since they employ the same mesoscale physics despite genetic divergence. Agent-based processes can lead to lineage-specific behaviors and morphological motifs, but also convergent or parallel ones if they act in analogous fashions. See main text for additional examples of generic and agent effects, and descriptions of their morphogenetic roles.

In the remainder of this section we will discuss two long-studied cases of such generic-agential synergy: (i) the formation and migration of multicellular slugs in dictyostelids, and (ii) formation of complex morphologies in fruiting bodies of both dictyostelids and myxobacterial species.

### Slug formation in Dictyostelium

When starvation drives *D. discoideum* into development the liquid-like streams that form culminate in aggregation centers. The mature aggregates, slugs, migrate over the surface in response to light and temperature gradients. Inside the slug, moving cells form smooth flow patterns similar to those of individual particles in liquids (Vasiev and Weijer, 2003). The slug is a long (~1 mm), thin (~50 μm) cylindrical mass with a well-defined anterior tip that directs its movement. During aggregation and early slug formation presumptive stalk and spore cells are sorted out along the anterior-posterior axis, and their relative positions become inverted in a ‘reverse fountain’ manner as the fruiting body forms.

This process exhibits both generic mesoscopic properties but also agent-like behaviors of the constituent cells. Odell and Bonner (1986), for example, used a continuum mechanics model of viscous flow in which cells moved both longitudinally, in response to an anterior-posterior cAMP gradient and transversely, in response to an unspecified gradient, to generate a rotational movement that could generate a rolling flow. Jiang et al. (1998) employed a discrete lattice model in which movement was determined by chemotaxis towards a center (the tip) and energetics (cell-cell adhesion), and found that with the right balance of the two forces, a reasonably correct pattern of sorting out resulted. Umeda and Inouye (2004) formulated a continuum model of a viscoelastic fluid made up of heterogeneous actively moving points (cells) that differed in various respects including their diffusive tendencies and abilities to offer resistance, and obtained, in addition to sorting out, plausible equilibrium shapes for the slug. Hogeweg, Marée, and coworkers combined agent-based and generic mechanisms – chemotaxis to cyclic AMP, differential adhesion and pressure generation - to simulate the aggregation of cells, the correct spatial distribution of cell type and their self-organization into a fruiting body (Marée and Hogeweg, 2001; Marée, 2000; Marée et al., 2013; Savill and Hogeweg, 1997). Trenchard (2019) has proposed a different agent-based mechanism for sorting, one that depends on differences in speeds of movement and energetics.

### Fruiting body branching

In contrast to *M. xanthus* and *D. discoideum* which exhibit branchless fruiting bodies, many of the species in both of their lineages develop into branched structures (Schaap et al., 2006; Yang and Higgs, 2014). In Dictyostelia, branches develop as the product of either budding or from a secondary cellular mass generated through pinching off of the main cellular mass (Schaap et al., 2006). These mechanisms can lead to different branching patterns in different species, with in some cases arrays of secondary fruiting bodies arranged about a primary axis of stalk cells (Gregg et al., 1996). In Myxobacteria, where evidence is more limited, branches seems to develop exclusively by budding of the main cellular mass; pinching off has not been reported in this group (Qualls et al., 1978). Also, regularity in the branch distribution, as observed for whorl-developing fruiting bodies in some Dictyostelia species, is not obvious.

Cox and co-workers have carried out detailed studies on the genesis of the branching pattern in fruiting bodies of the dictyostelid *Polysphondilium pallidum* (now *Heterostelium pallidum*, Sheikh et al. (2018)), and their studies point to the integrated functioning of generic and agent-like processes (reviewed in Bonner and Cox (1995). *P. pallidum/H. pallidum* fruiting bodies are the result of secondary cellular masses being pinched off in regular intervals from the primary cell mass as it moves upward as the main stalk is formed (Byrne and Cox, 1987). The secondary masses turn into whorls of regularly spaced branches perpendicular to the main stalk (McNally et al., 1987; McNally and Cox, 1988). As in *D. discoideum, P. pallidum/H. pallidum* elongation involves chemotactic movements towards a cAMP gradient, the source of which is a set of cells found at the tip of the cellular mass.

The mechanisms underlying pinching off of the secondary cellular masses remain unknown. However, since this takes place before branching, the cellular mass may still retain its liquid-like properties. Liquids may undergo pinch-off as a consequence of an imbalance of the velocities of individual subunits across the mass. If the velocities are sufficiently large, the adhesion forces will not be strong enough to keep the cellular subunits together and a (partial) pinch-off would occur. As with slug locomotion, described above, chemotaxis could induce a velocity gradient of the cells across the mass. Biased movement due to chemotaxis, along with the oscillatory intracellular dynamics, may help to explain the observed regularity in the spacing between the multiple secondary masses. This outcome, which is not trivially predicted from the generic behavior of the liquid-like primary mass, may thus depend on agent-like behavior.

The secondary cellular mass remains attached to the stalk and rounds up as expected for a liquid composed of homogeneously cohesive particles (McNally and Cox, 1988). Branches developed from the secondary mass are regularly arranged across the plane perpendicular to the main axis. The positions of the branches are proposed to be determined by a local activation-long range inhibition effect like that described by Turing (1952), although the components of this reaction-diffusion system have not been characterized (Cox et al., 1988).

The mechanism of branching itself is more problematic, since it is not an expected morphology of liquid-like materials. Plant tissues, however, routinely undergo budding and branching, an effect that has been attributed to the inherent properties of their material identity as deformable solids (Benítez et al., 2018; Hernández-Hernández et al., 2012). These motifs are independently recurrent developmental outcomes in all lineages of photosynthetic eukaryotes, including the various polyphyletic algal clades and the monophyletic land plant clade, the embryophytes (Hernández-Hernández et al., 2012). Since both Dictyostelia and Myxobacteria undergo solidification via ECM deposition and possibly liquid-to-solid jamming in portions of the multicellular mass after aggregation has been completed, this might allow the multicellular masses to escape from the physical constraints imposed by the liquid-like behavior and acquire the properties of deformable solids for which budding and branching are easily achievable.

In addition to the transition from a liquid-like behavior to a solid one, a differential increase of volume in the direction of the future branch is required to extrude from the main cellular mass a secondary mass that will bud and finally turn into a mature branch. In plants, this is achieved by localized cell proliferation in response to gradients of hormones (Benkova and Bielach, 2010; Vermeer and Geldner, 2015). In Myxobacteria and Dictyostelia, development proceeds with little, if any, cell division. One of two mechanisms, or a combination of them, might cause the required increment in volume: further deposition of ECM or expansion of individual cell volume. In either case, volume increase must occur in an irregular distribution over the mass, with foci of hyperplasia specifying the sites where branches will develop further.

While some myxobacterial species also have branched fruiting bodies (see, e.g., Zhang et al. (2003)), the lack of conventional chemotaxis (although see Taylor and Welch (2008) for a chemotaxis-like effect in these organisms) and molecular networks for local activation-long range inhibition may account for pinch-off and regular patterning in branching, respectively, not being observed during fruiting morphogenesis in Myxobacteria. It should be noted that fruiting bodies in these species grow vertically in a series of tiers, each involving the addition of a cell monolayer. The rate of formation of new tiers is too rapid to be attributed to cell division, which suggests that cells may be recruited from lower layers (Copenhagen et al., 2020; Curtis et al., 2007). This mechanism for vertical growth is robust in the face of diverse mutations and conditions, which suggest that it is an essential process in fruiting body morphogenesis (Curtis et al., 2007). Since it has been reported that the deposition of tiers can be slightly asymmetrical (Curtis et al., 2007), it is possible that branching in Myxobacteria arises from the amplification and robust reinstitution of such asymmetries across generations.

## DISCUSSION

Motivated by the parallelisms between the two major known lineages of multicellular aggregative organisms: the prokaryotic myxobacteria and the eukaryotic dictyostelids, we have reviewed the factors determining the main developmental events in these organisms. We suggest that as a consequence of cell-cell contact during aggregation, the nascent multicellular masses of each organism acquire liquid-like properties and thereby become subject to morphogenetic processes characteristic of such materials. This allows them to be studied, and in some respects explained, in terms of physical principles at the mesoscale. As expected from the physical theory, the cell aggregates can exhibit streaming, rippling, and rounding-up behaviors like those observed in non-living liquids.

While the molecules that mediate liquid-type properties in the two classes of organisms are largely different, the physical processes mobilized at the multicellular scale are generic and in that sense are the “same.” Furthermore, later in development cellular masses solidify and behave as deformable solids, another category of material with nonliving counterparts with generic properties. For such materials, branching is a predictable morphological outcome.

Although the behaviors in aggregating cells resemble those exhibited by non-living liquids, mathematical and computational models have also needed to include agent-based behaviors in addition to generic ones to achieve verisimilitude (Cotter et al., 2017; Fujimori et al., 2019; Janulevicius et al., 2015; Marée and Hogeweg, 2001). Unlike the molecular subunits of nonliving liquids, the cells constituting the multicellular masses can change and adapt their behaviors in response to external cues through complex regulatory and signaling pathways. We attribute the deviations of the dynamics and morphological outcomes of the multicellular masses from generic physical predictions to the contribution of agent-like behaviors, e.g., directed migration, regulated quiescence, oscillatory signal relay, reaction-diffusion coupling, of the cells themselves. Cells of clonally developing multicellular organisms can also exhibit agent-like behaviors (Christley et al., 2007; Giniunaite et al., 2020b; McLennan et al., 2020). While it is difficult to quantify the relative contributions that each class of phenomena makes to the respective developmental processes, considering the extent to which morphogenetic outcomes are predictable from generic physical considerations we suggest that morphogenesis of Myxobacteria and Dictyostelia is more dependent on agent-like behaviors than that of animals or plants. This is almost certainly a function of their aggregative nature.

Because of the relative indifference of generic processes to molecular variation (adhesion, for example, can be mediated by many different classes of proteins and glycans), the gene products that first mediated the production of a form or structure in a species’ earliest ancestors need not be the same one that is active in its present members. Consequently, the gene products that mobilize generic effects can differ widely in different classes of organisms (e.g., animals, plants, social amoebae and bacteria), and even in sister species, due to developmental system drift (True and Haag, 2001). In contrast, generic processes are part of the physical world, and therefore do not evolve per se, although the physics involved in a given lineage’s developmental routines can change over phylogeny (Newman, 2019a).

Many of the genes involved in generic processes in animal and plant lineages predated or accompanied the emergence of multicellularity. In those lineages, morphogenesis and pattern formation can be characterized in terms of the dynamical patterning modules (DPMs) that mobilize specific physical forces and physicochemical effects to produce the respective structural motifs (Newman, 2019b; Hernández-Hernández, 2012; Benítez et al., 2018). Similarly, some gene products that shape dictyostelids and myxobacteria as multicellular materials were carried over from single-celled ancestors, as were some gene products involved in agent behaviors. However, as we have described with the *M. xanthus* stringent response suppressive products CsgA and SocE, and the *D. discoideum* starvation-regulated paracrine factor Countin, some agent-associated genes seem to be novelties of the social forms.

While DPMs are, by definition intrinsically multicellular, agents are intrinsically individual – cellular, in the cases discussed here. Another important distinction is that agents are peculiar to the biological world, even if they are artifactual (e.g., robots). Thus, in contrast to generic materials, which have physically predictable macroscopic properties and behaviors, cellular agents have no such constraints on their activities. The rules they follow in developmental systems are as varied as cell behaviors (e.g., motility, secretion of ions, small and macro-molecules, electrical, chemical, and mechanical excitability) and responses to microenvironmental complexity permit.

Early comparisons between Myxobacteria and Dictyostelia noted that the morphological outcomes of their respective developmental processes resembled one another to a remarkable extent despite their independent origins, the evolutionary distance between them, and the lack of gene-based homology in the relevant mechanisms in the two groups. Our attention to this phenomenon was inspired by comparative analysis of the two lineages by Bonner (1982) and Kaiser (1986). Both favored explanations based on convergent selection for adaptation to similar ecological niches, with a focus on common developmental mechanisms such as cell adhesion, communication and oscillations (Kaiser, 1986) and “developmental constraints” such as that incurred by increased size (Bonner, 1982; Bonner, 2015). Based on the literature reviewed here, we conclude that the similar developmental trajectories and outcomes of Myxobacteria and Dictyostelia are more likely due to shared generic physical processes in coordination with analogous agent-type behaviors than to convergent evolution under parallel natural selection regimes. However, we acknowledge, in agreement with both Kaiser (1986) and Bonner (2015), that ecology, in the form of exploitation or construction of suitable environmental niches, is an essential factor in accounting for the establishment of these social phenotypes. Our analysis extends beyond the molecular mechanisms considered by these earlier investigators, to also include the physical nature of the multicellular masses. This approach is based on experimental and theoretical advances made in material sciences, particularly as applied to biological systems, in the intervening decades (see Forgacs and Newman (2005)), and progress in agent-based concepts and models (Thorne et al., 2007).

Some authors have noted the tendency of aggregative multicellular organisms to exhibit a narrower and simpler morphological diversity when compared to clonal organisms such as animals and plants (Grosberg and Strathmann, 2007). A common explanation to this observation is the emergence of genetic conflict arising between different cellular lineages being incorporated into the same conglomerate during aggregation. Despite kin selection mechanisms of “cheater” control (Travisano and Velicer, 2004), it is held that the impact of genetic conflict could still be large enough to destabilize multicellular structure and impair the evolution of further complexity. In clonal organisms, genetic conflict is thought to be avoided at every generation by genetic bottlenecks that reduce genetic diversity to those mutations emerging as consequence of DNA replication (Folse and Roughgarden, 2010). In his treatment of the evolution of Dictyostelia, Bonner (1982) also suggested that selective regimens are dependent on the scale on which they operate, and that size contributes to the differences in diversity between Dictyostelia and Myxobacteria compared with plants and animals.

The physical framework addressed here provides an alternative to the multilevel selection and scale-based accounts. As described above, despite the fact that animals, Dictyostelia and Myxobacteria can all be conceptualized as non-living liquids, the weaker associations between cells and surfaces in the social amoebae and bacteria lead to behaviors not observed in animals (e.g., streaming) and the stronger, cytoskeletally linked attachments in animals mediate behaviors (multilayering and lumen formation) not seen in the aggregative systems (Newman, 2019c). These differences are amplified by the fact that polarity (affecting, variously cell surface or shape in the different systems) is much more transient in Dictyostelia and Myxobacteria than in animals (Gómez-Santos et al., 2019; Manahan et al., 2004; Szadkowski et al., 2019), undermining the persistence of complex organization in the former two groups.

An important implication of the perspective we have presented here is that physics-based and agent-based approaches to understanding development are not simply alternative modeling or computational strategies, but represent realities of complex biological systems that are represented to various extents in different organismal lineages. Thus, the material nature of multicellular systems and the inherent structural motifs entailed by the relevant physics introduces a predictability to morphological evolution (Newman, 2016; Newman, 2019b). In contrast, agent-type behaviors are more unconstrained and open-ended in their possibilities, and their evolution could have led phylogenetic lineages that embody them (e.g., vertebrates, which have the novelty of a neural crest (York and McCauley (2020)) in less predictable directions.

Comparative analyses often rely on the study of homologous characters (i.e., those sharing common ancestry) in order to disentangle phylogenetic relationships and hypothesize evolutionary scenarios. These studies, mostly conducted in the population genetics framework underlying the evolutionary Modern Synthesis, have provided important insights regarding the processes of divergence of species as the product of selective pressures, genetic drift, mutation and gene flow (Pigliucci and Müller, 2010). But (with some exceptions, see Abouheif and Wray (2002)) they have generally neglected the role of development and, lacking a mechanistic view of phenotypic innovation (Müller and Newman, 2005), are limited in the extent to which homology can be assigned between characters in disparate groups (Müller, 2003; Müller, 2017).

Structures are considered homologous developmentally if they have the same form by virtue of having the same generative processes. Here we have invoked a more general sense of this concept, including in the notion of “sameness” generic physical mechanisms in addition to genes. In this we are echoing the insights of the Soviet biologist N.I. Vavilov, who in his classic paper “The law of homologous series in variation” wrote, “[g]enetical studies of the last decades have proved even the divisibility of the minutest morphological and physiological units in systematics…and established that, although outwardly similar, they can be different genotypically” (p. 48), and that “the great majority of varietal characters, not only within the limits of single genera and families but even in distant families, are homologous from a morphological point of view” (p 82) (Vavilov, 1922). We suggest that our broader concept of homology can help resolve enigmas of biological similarity across phylogenetic distances. Knowledge of molecular and cellular determinants of material identity and agent-like behaviors, in concert with suitable mathematical and computational models of these causally hybrid, multiscale systems (e.g., (Camley and Rappel, 2017; Cotter et al., 2017)), could ultimately provide a compelling and testable account of these morphological affinities.

## Acknowledgments

This paper is dedicated to the memory of our student and colleague, Juan Arias Del Angel. Juan framed the questions that led to this paper and wrote the first draft during a study visit to the SAN’s laboratory in the summer of 2019. Juan was supported by a Consejo Nacional de Ciencia y Tecnología (CONACYT) scholarship and was a recipient of a Company of Biologists (U.K.) traveling fellowship during his work on this project. His completed Ph.D. thesis in the Programa de Doctorado en Ciencias Biomédicas Universidad Nacional Autónoma de México, based on work cited herein, was submitted after his untimely death in November. We thank Sue Seif for drawing Figure 1 and Luis Guillermo García Jácome for drawing Figure 2.

